# Using multiscale molecular modeling to analyze possible NS2b-NS3 protease inhibitors from medicinal plants endemic to the Philippines

**DOI:** 10.1101/2023.05.12.540544

**Authors:** Allen Mathew F. Cordero, Arthur A. Gonzales

**Affiliations:** Department of Chemical Engineering, University of the Philippines Diliman, Quezon City, Philippines

## Abstract

Philippine folkloric medicinal plants like *Euphorbia hirta* (locally known as tawa-tawa), *Carica papaya* (papaya), *Psidium guajava* (guava), and *Momordica charantia* (bittermelon) have been used as a treatment for dengue. However, limited studies have been conducted regarding the extensive effects of these plants, especially their anti-dengue activity. This study evaluated 2,944 ligands from phytochemicals found in various medicinal plants as potential dengue inhibitors that could be developed into cost-effective and efficient therapeutic agents. SwissADME and Chembioserver online servers were used to conduct tests on absorption, distribution, metabolism, excretion, and toxicity (ADMET) for all ligands, resulting in 1,265 compounds being pharmacologically viable. By targeting the NS2b-NS3 protease of the dengue virus, specifically its catalytic triad of Asp 75, Ser 135, and His 51 residues, we can inhibit the replication of the virus. Molecular docking results showed ten ligands with comparable docking scores to the reference compounds. Attachment to the binding site is strengthened by electrostatic, polar, and hydrophobic interactions and the formation of hydrogen bonds.

Furthermore, we also evaluated their stability using molecular dynamics simulations on GROMACS 2021.3. Molecular dynamics simulations of up to 100 ns of chemical time suggest eight of the ten candidate ligands are stable while binding to the active site. Free energy calculations using molecular mechanics Poisson-Boltzmann surface area also proved that six of the eight stable ligands exceeded the binding energies of the reference compounds. Results showed that veramiline from *Veratrum mengtzeanum* (pimacao), etiolin from *Lilion martagon* (Turk’s cap lily), hydroxyverazine from *Eclipta prostrata* (false daisy), chlorogenin from *Yucca gloriosa* (palm lily), cyclobranol from *Euphorbia hirta* (tawa-tawa), and ecliptalbine from *Eclipta alba* maintained their structural stability throughout the simulations. They also displayed good oral bioavailability and potential drug-like characteristics. These six compounds warrant further *in vitro* and *in vivo* investigation as potential dengue therapies.

## Introduction

Over the past five decades, dengue virus (DENV) infections have shown a 30-fold increase globally, and 50 to 100 million new infections are estimated to occur annually in more than 100 endemic countries. In fact, around 2.5 billion people worldwide around at the risk of disease, with 20 million cases a year in more than 100 countries (1). In the Philippines, an alarming rate of 220,705 dengue virus type II (DENV-2) cases were recorded from January 1 to December 17, 2022, which is 182% higher than the 78,223 cases reported in 2021 (2).

Control strategies and intensive research against dengue virus (DENV) have become a global health priority to address the alarming rate of victims. Various studies have been conducted at the industrial and academic levels for years, but no clinical drugs that act directly against dengue virus (DENV) are currently available. However, one dengue vaccine has been approved, CYD-TDV, sold under the name Dengvaxia in 2015 (1). The vaccine was considered safe and expected to promote public health benefits when used in regions with approximately 70% seroprevalence in the age group eligible for the vaccine. Still, it should only be administered to people not previously affected by DENV (3).

According to a recent study (4) by Uday et al., it has been discovered that the known drug molecules idelalisib (IDE) and nintedanib (NIN) can effectively attach to the active site of NS2b-NS3 protease, an essential component for DENV replication process, leading to changes in the dynamics and conformation of the enzyme. In this protein, the active site is formed by the catalytic triad, composed of the amino acids His51, Asp75, and Ser135. These amino acids are in a specific configuration that enables them to act together as a nucleophile, a base, and an acid, respectively, during the cleavage of peptide bonds (5). Moreover, the triad is also responsible for recognizing and cleaving specific amino acid sequences within the polyprotein of DENV, thereby enabling the virus to replicate and propagate. The catalytic activity of the triad is critical for the function of the NS2b-NS3 protease and represents a potential target for developing antiviral drugs to combat dengue fever. Thus, the high binding energies of these drug molecules to the active site make it a good reference compound for the study.

In the Philippines, folkloric medicinal plants have been used as an alternative treatment against dengue fever because of their safety and cost-effectiveness. Approximately 30 different plant species have been found to have the potential to combat the virus. Among them are tawa-tawa (*Euphorbia hirta*), papaya (*Carica papaya*), guava (*Psidium guajava*), sambong (*Blumea balsamifera*), and bitter melon (*Momordica charantia*). The leaves of *E. hirta*, *P. guajava*, and *M. charantia*, are used to make a decoction that alleviates viral infection and associated fever symptoms (6). The study then selected and investigated 2,944 ligands from the phytochemicals of these medicinal plants to test their effectiveness against dengue.

The study primarily focused on targeting the NS2b-NS3 protease, as molecular dynamics (MD) simulations, virtual screening, and molecular docking were employed to find the protease’ best inhibitory binding energies, confirming the active site residues and their interactions and, ultimately, discover potential anti-dengue drug candidate. Herein, the interactions of small molecule inhibitors with the least binding energy were obtained and systematically analyzed. The binding site’s major and minor interaction points were demonstrated by the combination of ligand-based and structure-based approaches, providing a new strategy for designing novel and specific dengue virus therapeutic agents safe for human consumption.

Ultimately, the study warrants exploring the bioactive compounds and possible mechanisms of action and will pave the way to develop a new antiviral drug lead from *E. hirta* and other medicinal plants.

## Materials and methods

The crystallized 3D structure of NS2b-NS3 protease from the DENV-2 serotype was obtained from the Protein Data Bank (PDB) of the Research Collaboratory for Structural Bioinformatics (RCSB), PDB ID: 2FOM. Before docking experiments, protein crystal structures were prepared by removing ligands, adding necessary hydrogen atoms, optimizing hydrogen bonds, and removing solvents and metals/ions using Avogadro (7) and Schrodinger Maestro (8).

Structures of reference compounds idelalisib (IDE) and nintedanib (NIN) were downloaded from PubChem (www.pubchem.com) (9). Idelalisib and nintedanib are drug molecules used mainly in cancer treatment and were considered reference compounds based on the results of Uday et al. (2021) (4). On the other hand, ligand structures were obtained from Dr. Duke’s Phytochemical and Ethnobotanical database (10) and the comprehensive phytochemical database of the Molecular Modeling Research Laboratory of UP Diliman Department of Chemical Engineering. All structures were optimized using the energy minimization tool of Maestro (8) to maximize ligand structures for protein binding, improve the accuracy of docking results, and increase the efficiency of computational screening methods.

### Test for pharmacological viability

Biocompatibility refers to the ability of a substance or material to interact with living cells, tissues, or organs without causing harm or adverse reactions. Biocompatibility is crucial in drug development because drugs are designed to interact with specific targets in the human body. Any harmful effects on healthy cells or tissues can lead to serious side effects and health complications.

One way to ensure the biocompatibility of drug candidates is to test for their chemical absorption, distribution, metabolism, excretion, and toxicity (ADMET) using computational tools like SwissADME (www.swissadme.ch) (11) and Chembioserver (www.chembioserver.vi-seem.eu) (12). SwissADME (11) is a web-based platform that allows researchers to predict the pharmacokinetic properties of small molecules, such as drug candidates, based on their chemical structures. Here, Lipinski (13), Ghose (14), Veber (15), Egan (16), and Muegge (17) filters were applied, as they predict the drug-likeness of a compound with a specific biological activity designed for the oral route of administration. Lipinski filter recommends molecular weight (MW) of less than 500, lipophilicity (expressed as the logarithm of the partition coefficient between water and 1-octanol, log P) value of less than 4.15, and the number of O and H atoms not exceeding ten each. The Ghose filter defines drug-likeness constraints: calculated log P is between −0.4 and 5.6, MW is between 160 and 480, molar refractivity (MR) is between 40 and 130, and the total number of atoms is between 20 and 70 (14). Veber filter includes a less selective constraint such as rotatable bonds (RB) of less than ten and total polar surface area (TPSA) of less than 140 Å while Egan recommends log P of less than 5.88 and TPSA of less than 131.8 Å (15,16). For Muegge’s rule, the constraints are MW between 200 and 600, log P of −2 to 5, TPSA of less than 150 Å, number of rings less than 7, at least four carbon atoms, number of RB less than 15, hydrogen bond acceptor (HBA) less than 10, hydrogen bond donor (HBD) less than 5 (17).

The structures of the ligands were also uploaded onto ChemBioServer (12), which can detect certain toxic functional groups and evaluate their toxicity.

### Molecular docking

Molecular docking was carried out using Autodock 4.2 (18), implementing a Lamarckian Genetic Algorithm (LGA). Autodock Tools 1.5.6 generated a grid box surrounding the selected residues: His51, Ser135, and Asp75. The grid box also corresponds to the location of the active site, which was determined to be the catalytic triad of the three selected residues (19,20). The grid center coordinates for the binding cavity of NS2b-NS3 protease (2FOM) were set to −2.249, −9.192, 16.146 in the x, y, and z direction, respectively, with a grid box size of 50 x 50 x 50 Å (0.375 Å spacing). For exhaustiveness, the number of GA runs was set to 1000, with 1000 poses generated per run. From this, only the best conformation (one with the most negative binding energy) per run was selected.

Each complex’s best docking conformations were compared through their binding energies, in which the ten ligands with the highest binding energy were selected. Autodock output (.pdbqt) files were converted to .pdb format through Schrodinger Maestro (8). The same software was also employed to analyze the interacting residues from NS2b-NS3 protease.

### Molecular dynamics simulation

Molecular dynamics (MD) simulation evaluates the conformational stability of the complex and predicts any changes in the structure within a specific period. The best conformation of protease and protease-ligand complex systems were subjected to MD simulations with the help of the Groningen Machine for Chemical Simulations (GROMACS) 2021.3 (21). For the protein, the Chemistry at Harvard Macromolecular Mechanics (CHARMM) 36 force field - July 2022 parameter was used throughout the whole simulation process. On the other hand, the ligand’s force field parameters were generated from the CGenFF (https://cgenff.umaryland.edu/) server of the University of Maryland Baltimore (22). The ligand-protein complexes were dissolved in TIP3P water, containing 0.15 M NaCl ions (NaCl concentration in human blood). Molecular dynamics simulations of up to 100 ns of chemical time were performed, with a working temperature of 300 K and a constant pressure of 1 bar (0.9869 atm). Snots of the trajectory were saved every 100 ps.

### Molecular Mechanics/ Poisson Boltzmann Surface Area (MM/PBSA)

The Molecular Mechanics/Poisson Boltzmann (MM/PB) method provides a more accurate calculation of the binding strength between ligands compared to docking experiments. In the Molecular Mechanics/Poisson Boltzmann (MM/PB) approach, the overall energy of a molecule is calculated as the sum of its molecular mechanics’ energy (which considers van der Waals and electrostatic interactions), its polar solvation free energy, and an adjustment for the molecule’s configurational entropy (23). It can be expressed as follows:

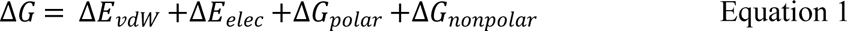

Non-polar solvation energy (ΔG_nonpolar_) can also be estimated by using the most widely used non-polar model, solvent accessible surface area (SASA) model. It includes the work done to create a cavity in the solvent and the attractive forces between the solvent and solute (24).

The MM/PBSA method was implemented through the g_mmpbsa tool (24) of GROMACS, with the help of the Python script MmPbSaStat.py. However, the tool cannot calculate the absolute binding energy due to a lack of entropic energy calculations (24).

## Results and Discussion

### Biocompatibility and toxicity test results

Phytochemicals or bioactive compounds found in plants are an alternative source of drug candidates, offering a vast array of natural molecules with diverse structures that have yet to be fully explored. In fact, 80% of the population in developing countries continues to rely on herbal extracts due to their demonstrated safety and effectiveness in traditional medicine (29). Generally, ligands from the database are aromatic hydrocarbons with high molecular weights (MW). Other compounds belonging to various chemical classes, such as flavonoids, quercetins, and natural chalcone (a type of natural ketone) compounds, agree with other studies.

Table 1 summarizes the results of chemical absorption, distribution, metabolism, excretion, and toxicity (ADMET) tests of 2,944 ligands using SwissADME (11) and Chembioserver (12). To assess the drug-likeness of each ligand, we counted the number of violations on five different filters (Lipinski, Ghose, Veber, Egan, and Muegge) and recorded them in S1 Table.

**Table 1.** Summary of biocompatibility and toxicity results for the test ligands.

| Source | No. of Ligands | No. of Ligands that Passed SwissADME | No. of Ligands that Passed ChemBioserver 2.0 |
| --- | --- | --- | --- |
| Comprehensive Phytochemical Database | 2849 | 1677 | 1200 |
| Dr Duke's<br>Phytochemical<br>Database | 95 | 88 | 65 |
| Total | 2944 | 1765 | 1265 |

Any compounds that had a total of five or more violations, as recommended by SwissADME for an oral drug (25), across all filters were deemed “failed” and were not subjected to toxicity testing. Imposing rigorous screening measures on the compounds is essential to understand their drug-like properties thoroughly. As such, Table 1 indicates that out of the 2,944 test ligands, only 1,765 have met the criteria set by SwissADME.

Among the five filters, the Muegge filter has given the most number of violations (3,456), closely followed by the Ghose filter (3,251). Both filters include the lipophilicity parameter (log P), mainly contributing to the ligand violations. Most phytochemicals have a log P value of less than five because they are highly non-polar. Among the parameters measured, it is possibly the least important physicochemical property of a potential drug since most recently discovered and synthesized drug molecules have high log P values (26). When absorbed by the body, the high stability of phytochemicals in gastric and colonic pH also helps absorption and bioavailability (27). While π-stacking interaction can increase the binding affinity of the inhibitor to its target, de Freitas & Schapira (2017) (28) suggests that reducing the number of aromatic rings of a molecule might improve its physicochemical properties, such as solubility.

Further screening of the ligands resulted in 1,265 ligands passing the toxicity test on Chembioserver. Table on S1 Table shows that failing ligands mostly contain catechol and butanone-Michael acceptors, which pose risks to human bodies. Catechol is an irritant to the skin, eyes, and respiratory system. Prolonged or repeated exposure to high catechol concentrations can cause skin sensitization and respiratory irritation and may increase cancer risk. On the other hand, limited information is available on the specific toxicity of butenone-Michael acceptor to humans. However, based on the chemical structure of this compound, it is likely to have the potential to cause harm to human health, as it is a reactive compound that can bind covalently to biological molecules such as proteins and DNA, leading to cellular damage and dysfunction. This chemical reaction can disrupt normal cellular processes, potentially leading to adverse health effects (12).

### Docking results

Table on S2 Table shows docking results for all 1,265 ligands, including the reference compounds idelalisib and nintedanib. All ligands came from plants, with some traditionally used as medicine in other parts of the world, including papaya, bitter melon, guava, basil, neem, cat’s claw, ginger, and lemongrass. The negative binding energies for all these ligands indicate favorable binding in all complexes, which concluded that the protein’s binding site was correctly found and bounded by the ligands. Among the test ligands, 22 (1.74%) have higher binding energy compared to the reference ligands idelalisib (−25.522 kJ/mol), while 20 (1.58%) exceeded nintedanib (−29.790 kJ/mol). In addition, they also have a higher molecular weight, resulting in more possible binding sites and interactions with the protein residues. In fact, according to a study (1), the molecular surface area of drug-like molecules increases by 95 Å for every 100-unit increase in molecular weight. Thus, higher molecular weight and lipophilicity rather than polar interactions drove most of the potent hits.

Fig 1 shows the best docking conformations of the ligands with the highest binding affinity to NS2b-NS3 protease, including the interacting residues. We can also observe that high docking scores came from their direct attachment to the protein. In addition, we can quickly notice the cleavage formed from the active site location (Fig 1), as it is the most active domain of the protein (1).

**Fig 1.**
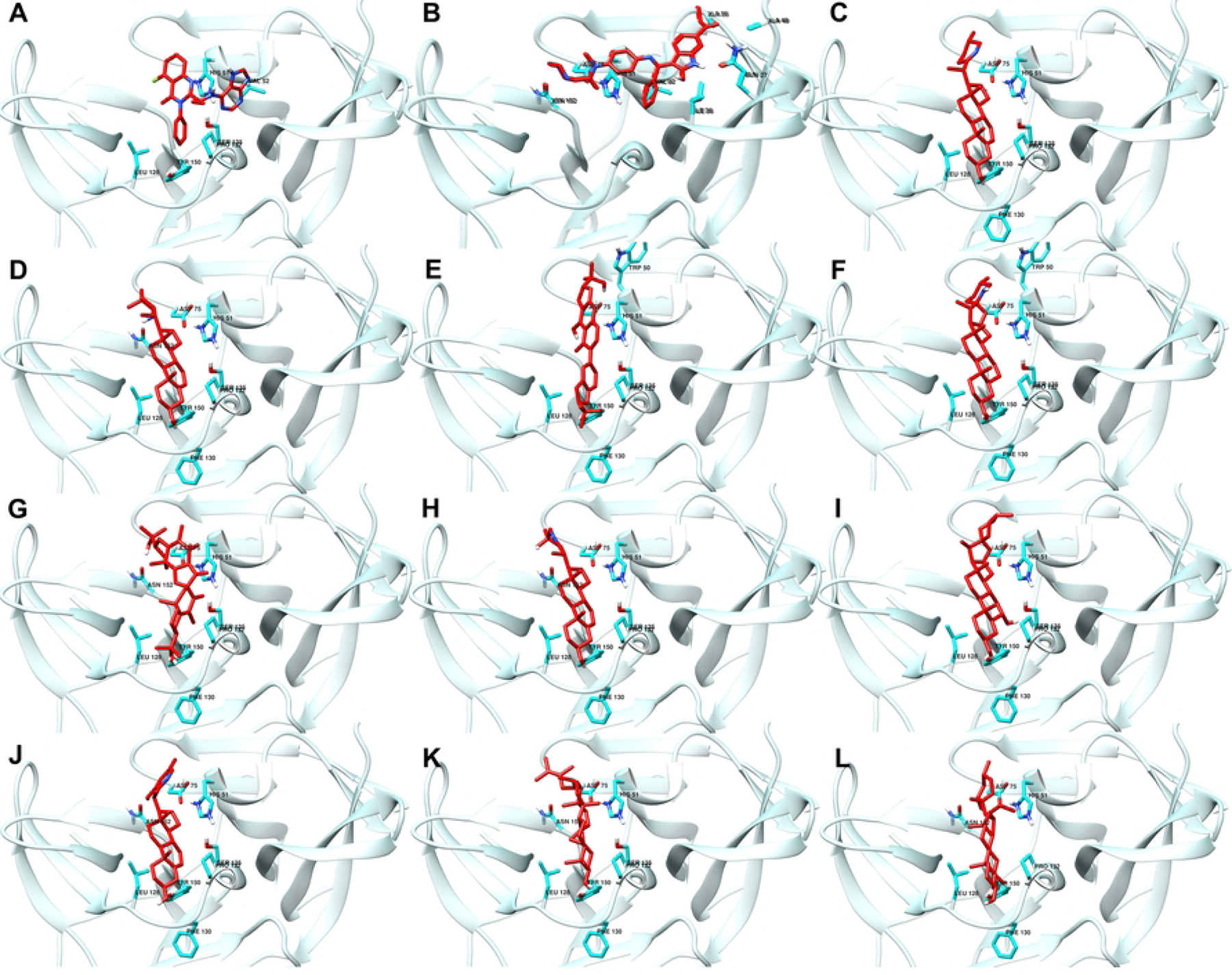
Best docking conformations of top ten ligand-protein complex based on binding energy. Reference ligands: (A) IDE (B) NIN. Candidate ligands: (C) VER (D) ISO (E) ETI (F) TOM (G) CAR (H) HYD (I) CHL (J) ECL (K) CYC (L) HON

Table 2 shows that CAR (−42.773 kJ/mol) and HON (−49.984 kJ/mol) are only the top ten ligands with lower MM/PBSA scores than NIN. We can also observe that the MM/PBSA energies do not follow the same trend as the binding energy from docking. For example, CYC has a higher MM/PBSA score (−70.943 kJ/mol) compared to CAR (−42.773 kJ/mol), despite having less binding energy (−34.309 kJ/mol to −36.987 kJ/mol respectively) from docking. The difference can be attributed to adding polar and non-polar solvation energies (Equation 1) in the MM/PBSA method calculation, resulting in more negative binding energy (30). Since all candidate ligands have a stronger binding affinity to the protein than the reference compounds, all of them were included in the succeeding experiments.

**Table 2.** Binding energy and MM/PBSA scores after molecular docking.

| Database no. | 3-Letter Symbol | Ligand | Plant source | MW (g/mol) | Docking score (kJ/mol) | MM/PBSA score (kJ/mol) |
| --- | --- | --- | --- | --- | --- | --- |
| Reference | IDE | idelalisib | - | 415.4 | -25.522 | -32.531 |
| Reference | NIN | nintedanib | - | 539.6 | -29.790 | -55.709 |
| 15559023 | VER | veramiline | pimacao | 509.6 | -38.953 | -80.682 |
| 10526985 | ISO | isolupinisoflavone | banyan fig | 438.5 | -38.786 | -20.628 |
| 168872 | ETI | etiolin | Turk's cap lily | 413.6 | -38.242 | -59.923 |
| 65576 | TOM | tomatidine | tomato | 415.7 | -37.865 | -15.984 |
| 101316738 | CAR | carindone | Carandas plum | 512.7 | -36.987 | -42.773 |
| 10549683 | HYD | 25beta-Hydroxyverazine | false daisy | 413.6 | -35.815 | -61.951 |
| 12303065 | CHL | chlorogenin | palm lily | 432.6 | -34.936 | -63.279 |
| 10692897 | ECL | ecliptalbine | false daisy | 409.6 | -34.727 | -56.932 |
| 39 | CYC | cyclobranol | Tawa-tawa | 440.7 | -34.309 | -70.943 |
| 52931465 | HON | hongguanggenin | century plant | 464.7 | -33.012 | -49.984 |

### Interacting residues

Most of the residues in the binding site of NS2b-NS3 protease that interacted with the two reference compounds also interacted with the ten candidate ligands, shown in Table 3. It contributes to the higher binding energy of the ten complexes when compared to the reference compounds (31). Active site residues Ser 135, Asp 75, and His 51 showed the most interactions, which also agrees with other studies (19,20,31–33). It is because the three residues also constitute the residues needed for the normal enzymatic function of the protein (31). Other interactions like polar and charged/noncharged (electrostatic) particle interaction can also affect the stability of the complex.

**Table 3.**
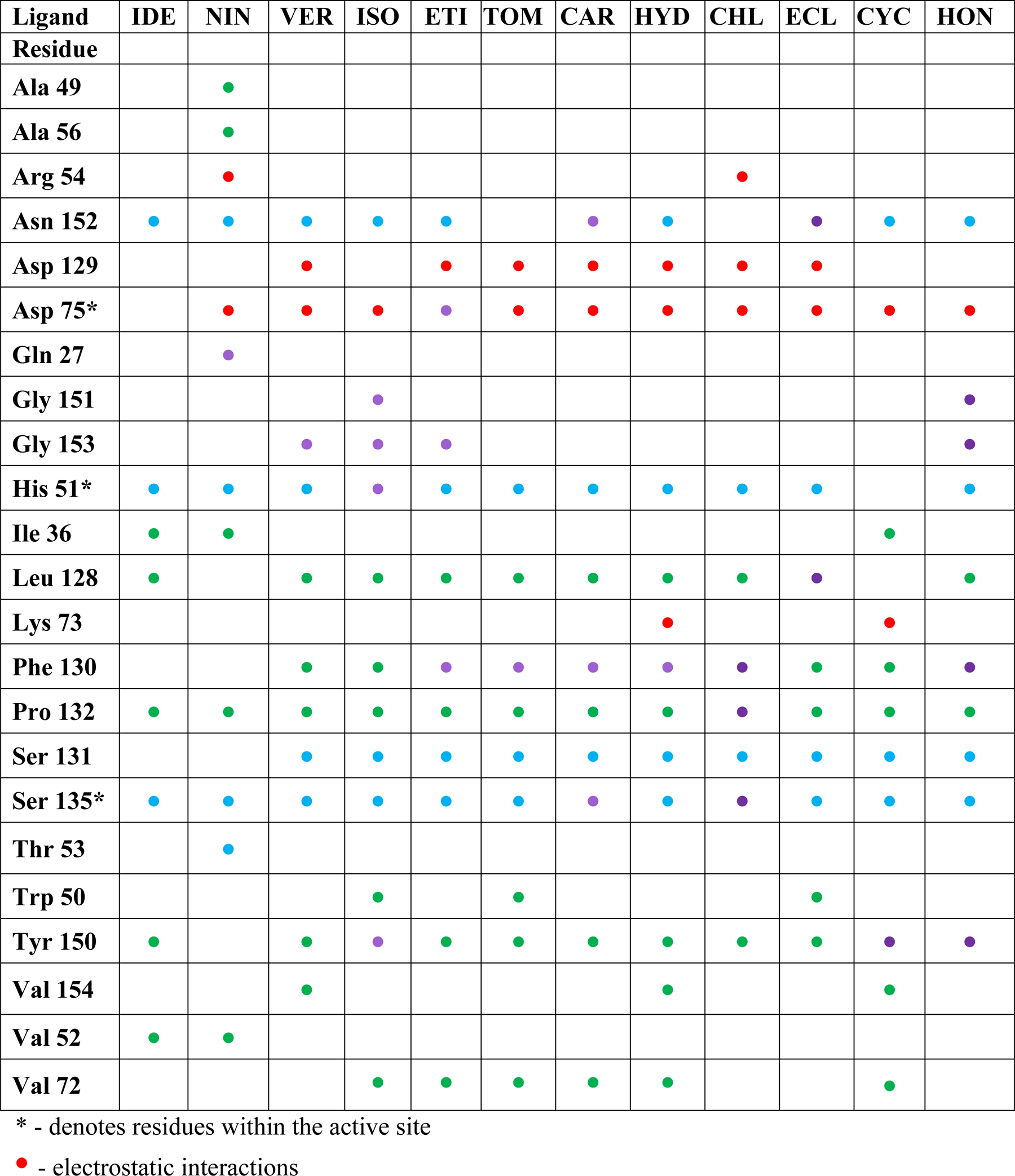
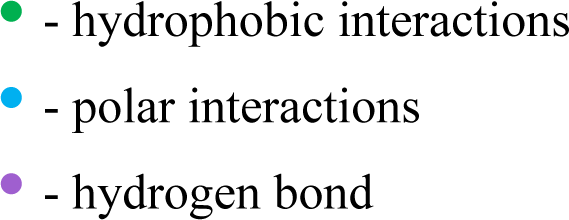
Summary of interactions between protein residues and references/ top ten ligands.

| Ligand | IDE | NIN | VER | ISO | ETI | TOM | CAR | HYD | CHL | ECL | CYC | HON |
| --- | --- | --- | --- | --- | --- | --- | --- | --- | --- | --- | --- | --- |
| Residue |  |  |  |  |  |  |  |  |  |  |  |  |
| Ala 49 |  | ● |  |  |  |  |  |  |  |  |  |  |
| Ala 56 |  | ● |  |  |  |  |  |  |  |  |  |  |
| Arg 54 |  | ● |  |  |  |  |  |  | ● |  |  |  |
| Asn 152 | ● | ● | ● | ● | ● |  | ● | ● |  | ● | ● | ● |
| Asp 129 |  |  | ● |  | ● | ● | ● | ● | ● | ● |  |  |
| Asp 75* |  | ● | ● | ● | ● | ● | ● | ● | ● | ● | ● | ● |
| Gln 27 |  | ● |  |  |  |  |  |  |  |  |  |  |
| Gly 151 |  |  |  | ● |  |  |  |  |  |  |  | ● |
| Gly 153 |  |  | ● | ● | ● |  |  |  |  |  |  | ● |
| His 51* | ● | ● | ● | ● | ● | ● | ● | ● | ● | ● |  | ● |
| Ile 36 | ● | ● |  |  |  |  |  |  |  |  | ● |  |
| Leu 128 | ● |  | ● | ● | ● | ● | ● | ● | ● | ● |  | ● |
| Lys 73 |  |  |  |  |  |  |  | ● |  |  | ● |  |
| Phe 130 |  |  | ● | ● | ● | ● | ● | ● | ● | ● | ● | ● |
| Pro 132 | ● | ● | ● | ● | ● | ● | ● | ● | ● | ● | ● | ● |
| Ser 131 |  |  | ● | ● | ● | ● | ● | ● | ● | ● | ● | ● |
| Ser 135* | ● | ● | ● | ● | ● | ● | ● | ● | ● | ● | ● | ● |
| Thr 53 |  | ● |  |  |  |  |  |  |  |  |  |  |
| Trp 50 |  |  |  | ● |  | ● |  |  |  | ● |  |  |
| Tyr 150 | ● |  | ● | ● | ● | ● | ● | ● | ● | ● | ● | ● |
| Val 154 |  |  | ● |  |  |  |  | ● |  |  | ● |  |
| Val 52 | ● | ● |  |  |  |  |  |  |  |  |  |  |
| Val 72 |  |  |  | ● | ● | ● | ● | ● |  |  | ● |  |

\* - denotes residues within the active site
● - electrostatic interactions
● - hydrophobic interactions
● - polar interactions
● - hydrogen bond

Most interactions are hydrophobic and polar, with a few electrostatic interactions (Asp 129 and Asp 75) present. Hydrophobic interactions are the most common type of interaction in protein-ligand complexes and usually determine the overall stability of the complex (28). As such, they are the main driving force in ligand-protein interactions. Most ligands formed hydrophobic contact with the amino acids: Leu 128, Phe 130, Pro 132, Tyr 150, and Val 72. Here, Phe 130 and Pro 132 hydrophobic interactions are present in all top ten ligands. Most hydrophobic interactions are created by an aliphatic carbon in the receptor (2FOM) and an aromatic carbon in the ligand. Since most ligands tested have aromatic carbons, we have similar observations with other studies (28,34) proving aromatic rings are prevalent in small-molecule inhibitors. In fact, 76% of the marketed drugs contain one or more aromatic rings, with benzene being the most frequently encountered ring system (34).

Hydrogen bonds (H-bond) are the prevailing directional intermolecular interactions in biological complexes and the predominant contribution to the specificity of molecular recognition (35). It is the main reason hydrogen bonds are exploited to gain specificity owing to their strict distance and geometric constraints in drug design(28). In this study, all top ten candidate ligands have at least one H-bond interaction - ISO has the most with four, while VER, TOM, and CYC have one. We can also observe that NS2b-NS3 residues were more often hydrogen-bond donors than acceptors (Fig 2). Glycine and phenylalanine are the most frequent H-bond acceptors due to the absence of a side chain to mask backbone atoms and increased backbone flexibility to satisfy the spatial constraints of H-bonds better (28). Some ligands (ISO, ETI, CAR, and CHL) also formed H-bonds with the active site residues His 51, Asp 75, and Ser 135. H-bond contacts were also present on other residues, including Leu 128 and Pro 132. The presence of hydrogen bonding and a higher number of hydrophobic and polar residues interactions could explain why the binding energy values of the candidate ligands are relatively higher than references IDE and NIN.

**Fig 2.**
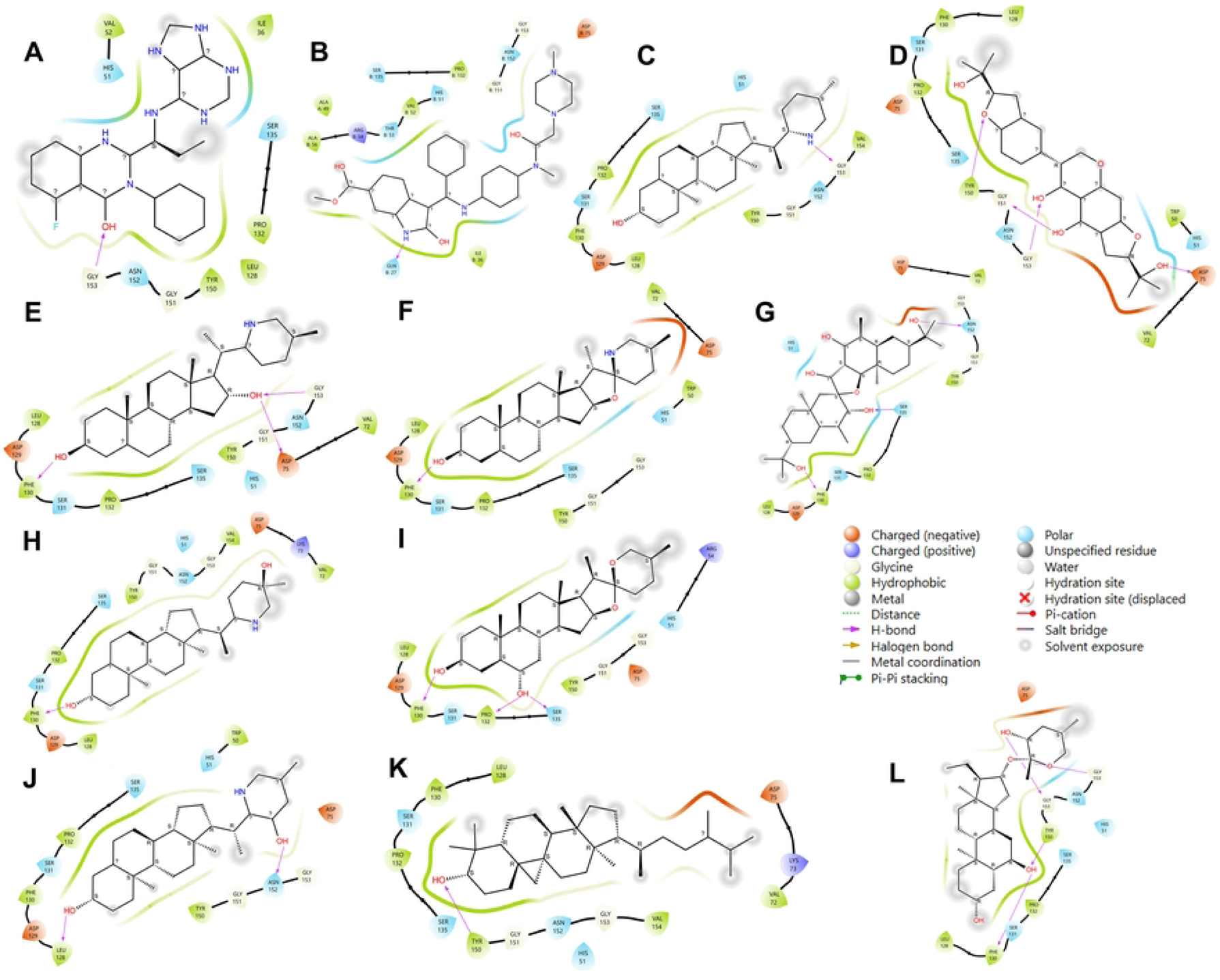
Illustration of interacting residues with the top ten ligands and references. Reference ligands: (A) IDE (B) NIN. Candidate ligands: (C) VER (D) ISO (E) ETI (F) TOM (G) CAR (H) HYD (I) CHL (J) ECL (K) CYC (L) HON.

### Root Mean Square Deviation (RMSD) analysis

For this study, the reference structure would be the best ligand-protease complex conformation obtained from molecular docking. In addition, systems no longer needed to be equilibrated for a time interval since the ligand structures docked were already minimized, and the reference structure already had the highest binding energy (36). Thus, the production run of 100 ns already provides the necessary trajectories for the RMSD calculation, with protein-ligand complex as the point of reference and backbone for least squares fit. Plots from Fig 3 illustrate the RMSD trajectories for the reference and top ten ligands.

**Fig 3.**
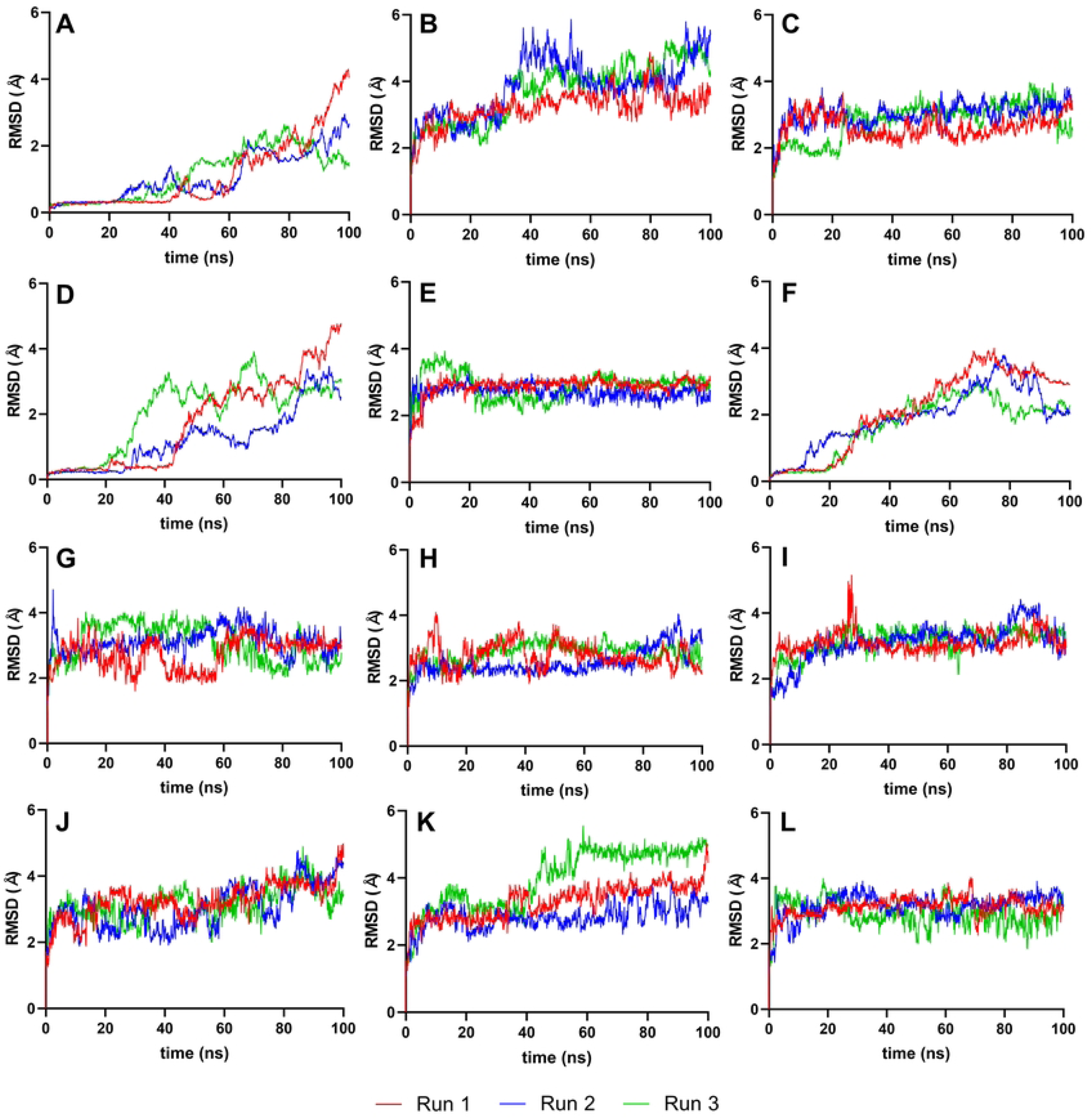
Complex-RMSD of the top ten complexes and two references at 100 ns. Reference ligands: (A) IDE (B) NIN. Candidate ligands: (C) VER (D) ISO (E) ETI (F) TOM (G) CAR (H) HYD (I) CHL (J) ECL (K) CYC (L) HON

Most ligand systems (VER, ETI, CAR, HYD, CHL, ECL, CYC, AND HON) have a steady RMSD trajectory behavior with a small average deviation (Table 4), an indication of high stability. However, ISO and TOM gradually increase RMSD over the simulation due to the systems still reaching equilibrium. Huge fluctuations at some points of the plot were also present, like a sudden rise in deviation of protein structure at 20-25 ns for ISO and 60 ns for TOM. It demonstrated significant instability, as several conformation changes and the highest RMSD values were observed in these systems. Table 4 shows the statistical analysis for the exact RMSD values obtained from each MD simulation.

**Table 4.** Statistical analysis of the RMSD values from the last 10 ns of MD simulation of top ten complex and reference ligands

| Ligand | Complex-Ligand RMSD (Å) |  |  |  | Ligand Only RMSD (Å) |  |  |  |
| --- | --- | --- | --- | --- | --- | --- | --- | --- |
|  | Mean | Min | Max | Standard Deviation | Mean | Min | Max | Standard Deviation |
| IDE | 1.5552 | 1.2403 | 1.8803 | 0.1416 | 0.1679 | 0.1528 | 0.1819 | 0.0550 |
| NIN | 4.8220 | 4.1350 | 5.1550 | 0.2290 | 2.8230 | 2.5560 | 3.2810 | 0.1340 |
| VER | 3.0030 | 2.1370 | 3.9440 | 0.4990 | 0.8400 | 0.6200 | 1.2670 | 0.1170 |
| ISO | 2.9716 | 2.4307 | 3.4640 | 0.2261 | 0.2649 | 0.1395 | 0.3073 | 0.3490 |
| ETI | 2.9840 | 2.5860 | 3.2530 | 0.1190 | 0.9060 | 0.5880 | 1.4130 | 0.1610 |
| TOM | 2.2119 | 1.9960 | 2.4587 | 0.1077 | 0.0682 | 0.0357 | 0.0967 | 0.1240 |
| CAR | 3.0390 | 2.7610 | 3.5880 | 0.1450 | 1.1630 | 0.8000 | 1.5620 | 0.1630 |
| HYD | 2.7650 | 2.3320 | 3.2450 | 0.2180 | 1.0940 | 0.6080 | 1.5640 | 0.2850 |
| CHL | 3.3190 | 2.7430 | 3.9370 | 0.3020 | 0.3830 | 0.2820 | 0.6720 | 0.0750 |
| ECL | 4.0040 | 3.4060 | 4.5590 | 0.2800 | 2.0320 | 1.6580 | 2.4090 | 0.1470 |
| CYC | 4.8640 | 4.3180 | 5.2050 | 0.1650 | 1.6270 | 1.1550 | 2.1040 | 0.2050 |
| HON | 2.8660 | 2.1630 | 3.5620 | 0.2990 | 1.0940 | 0.6080 | 1.5640 | 0.2850 |

Except for ISO and TOM complexes and the reference IDE, all other complexes reached equilibrium within 100 ns of simulation time with an RMSD range of 2-3 Å and 4-5 Å. From Fig 3, VER, ETI, CAR, HYD, CHL, and HON have stabilized fastest at 5-10 ns, whereas CYC and HON reached equilibrium at approximately 60 ns. However, during their initial MD runs, VER, CAR, and HYD took longer to equilibrate, about 60 ns.

Generally, a lower RMSD value with respect to the reference structure is good, as it represents good reproduction of the correct pose over some time (36). In line with this, smaller range and standard deviation of RMSD values indicate higher stability of molecules, as illustrated by VER, ETI, HYD, CHL, ECL, and CYC, which are all significantly better than the reference ligands, DAB and IDE.

Moreover, it is also important to show the RMSD trajectory of the ligands with respect to their best docking pose generated by AUTODOCK 4.2 to show high conformational stability (37). Fig 4 illustrates the RMSD trajectories, using ligands only as a reference point and least squares fit. The results imply that the ligands have only small movements inside the binding site, reaching as small as ∼0.5 Å, making them highly stable and reliable (36).

**Fig 4.**
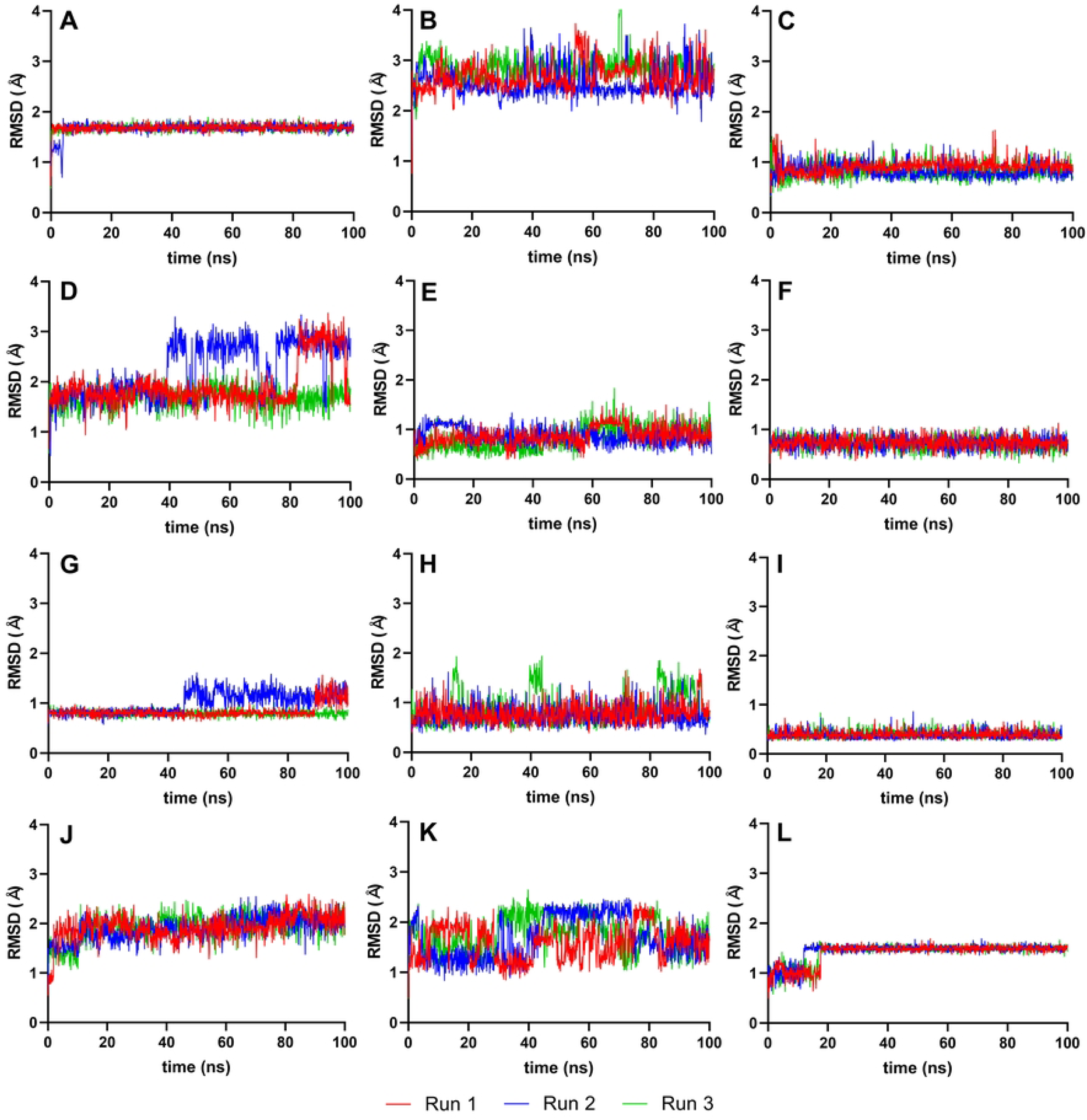
RMSD graphs for the reference and candidate ligands using ligands only as the reference point (Ligand-RMSD). Reference ligands: (A) IDE (B) NIN. Candidate ligands: (C) VER (D) ISO (E) ETI (F) TOM (G) CAR (H) HYD (I) CHL (J) ECL (K) CYC (L) HON

Like their complex-RMSD, IDE, ISO, and TOM also generated erratic and large fluctuations in their ligand-RMSD. These ligands’ large and more complicated structures may be attributed to their other interactions outside the protein’s binding pocket.

### Root Mean Square Fluctuation (RMSF) analysis

As the structures studied by MD simulations become more complicated, it becomes increasingly common to find high RMSDs related to the large fluctuations of structural subsets that do not reflect the structural changes of the macromolecule. Therefore, calculating the RMSF of protein is essential to help discriminate the flexible from rigid structures.

RMSF plots can also indicate the flexibility of each amino acid residue during molecular dynamics simulations. As seen in Fig 5, the RMSF values of the residues making up the active site of NS2b-NS3 protease showed minimal alteration. The troughs’ low values suggest that the residues had restricted movements, which prevented significant structural changes in the binding site and made it less susceptible to rearrangement. Since the site’s shape remained unchanged during the simulation, ligands could stay inside the cavity and attach themselves completely.

**Fig 5.**
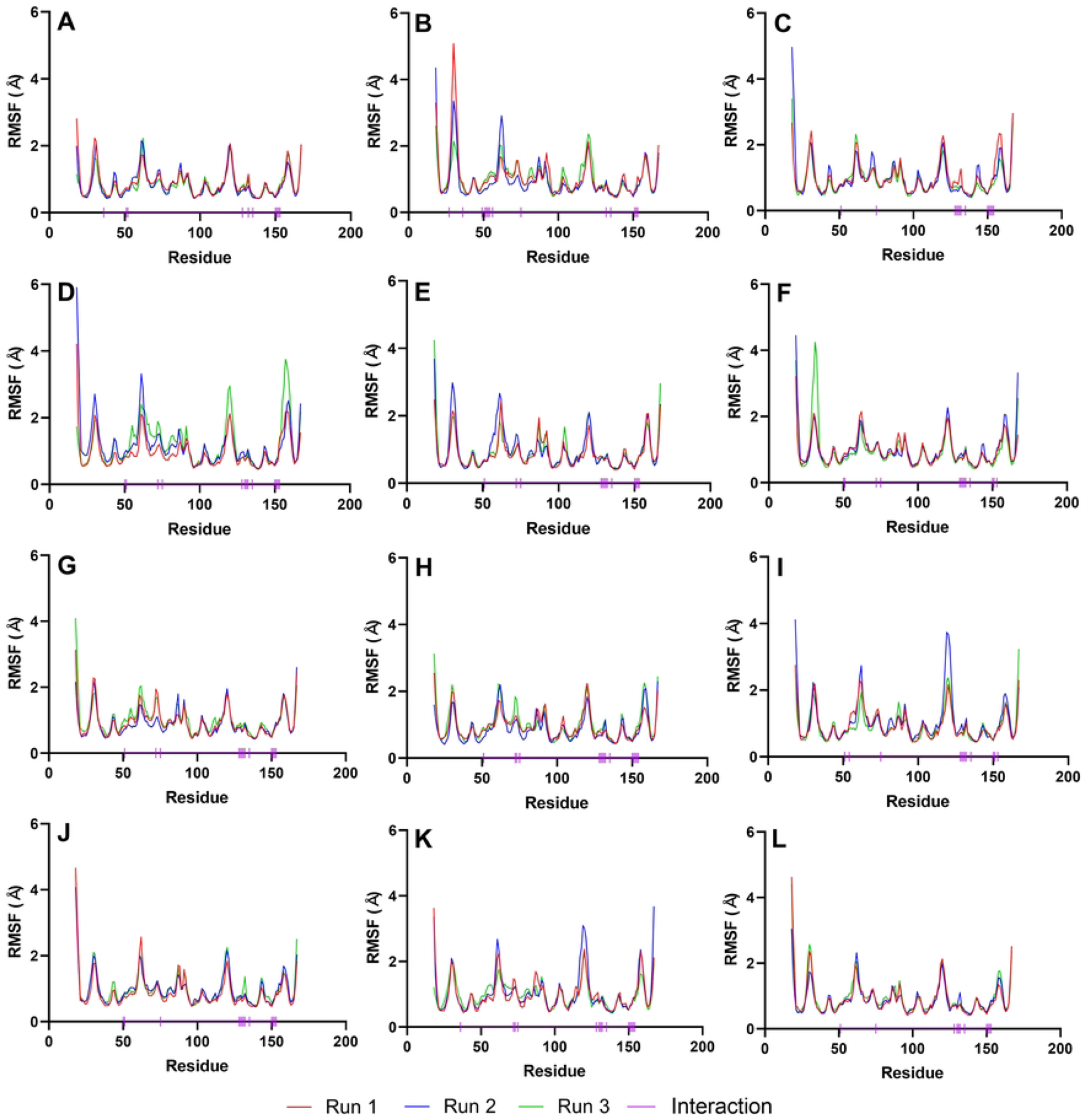
RMSF graphs of NS2b-NS3 Protease docked with the reference and candidate ligands. Residues with interactions were denoted by a small vertical line (|) on the x-axis. Reference ligands: (A) IDE (B) NIN. Candidate ligands: (C) VER (D) ISO (E) ETI (F) TOM (G) CAR (H) HYD (I) CHL (J) ECL (K) CYC (L) HON

From Fig 5, the RMSF for the loop regions is larger when the residues form the helix and sheet conformations from the ligand interactions, indicating that the secondary structures make the protein rigid. In addition, the protease (2FOM) has almost similar RMSF trajectories for most of the ligands, implying that they have the same formed interactions with the residues. The purple bars on the x-axis represent the position of the critical residues, including active site residues Asp 75, Ser 135, and His 51, and the gate residues Ser 131, Phe 130, and Tyr 150. As seen in Fig 5, all these residues coincide with the low RMSF values. The ligands that sufficiently formed contacts with the active site residues also had constrained movements.

### Molecular Mechanics Poisson-Boltzmann Surface Area (MM/PBSA) analysis

The binding affinities of the top ten ligands and reference complexes were determined using MM/PBSA binding free energy analysis. Only the part in equilibrium (final 10 ns) of their MD simulation trajectories was considered here.

The results show that six ligands (VER, ETI, TOM, HYD, CHL, and CYC) have higher MM/PBSA binding energy than the reference, NIN (−55.709 kJ/mol). VER still sits on top with −80.682 kJ/mol, followed by CYC with −69.763 kJ/mol. Despite having higher binding affinity from docking studies, CAR and HON have lower MM/PBSA binding energies when compared to NIN. Unstable reference (IDE) and candidate ligands (ISO and ECL) complex MM/PBSA energies were also not determined.

Molecules with large dipole moments and high polar surface area require more energy in the desolvation process, which would explain why ETI and HYD have high polar solvation energy (23). On the other hand, NIN, VER, and CHL only required relatively low solvation energies in the coupling process due to their lower number of polar interactions (Table 3). Since most ligands have similar structures (sterols, aromatic), we also expected their van der Waals or electrostatic energy to be close.

High dipole moment and electrostatic interactions (negatively/positively charged residues) also be attributed to the ligand’s electrostatic energy. ETI and HYD have the highest dipole moment among the ten candidate ligands, resulting in increased negative electrostatic energies. Table 3 also shows that HYD has the most positive/negatively charged interactions with the protein, contributing to the higher electrostatic energy.

In aromatic ligand-protein systems, the bonding and structure are predominantly determined by a complex interplay between van der Waals (vdW) and electrostatic interactions. Despite having lower electrostatic energies (Table 5), VER, CYC and HON have considerably high van der Waals (vdW, also called dispersion) energies, contributing to their high binding energy. When all energy terms were added, the VER and CYC could still surpass the binding energy of reference NIN. The other seven candidate ligands also have relatively strong van der Waals interactions, with energy values close to the ligands mentioned above.

**Table 5.** MM/PBSA binding energies of stable candidate ligands and reference.

| **Ligand** | **Van der Waals energy (kJ/mol)** | **Electrostatic energy (kJ/mol)** | **Polar solvation energy (kJ/mol)** | **SASA (kJ/mol)** | **Binding energy\* (kJ/mol)** |
| --- | --- | --- | --- | --- | --- |
| NIN | -78.447 | -1.298 | 34.100 | -10.063 | -55.709 |
| VER | -104.94 | -6.494 | 43.497 | -13.195 | -80.682 |
| ETI | -107.656 | -32.462 | 94.027 | -13.832 | -59.923 |
| CAR | -80.502 | -2.056 | 50.939 | -11.154 | -42.773 |
| HYD | -103.191 | -27.037 | 81.377 | -13.100 | -61.951 |
| CHL | -82.811 | -1.823 | 31.861 | -10.506 | -63.279 |
| ECL | -76.444 | -9.489 | 39.667 | -10.666 | -56.932 |
| CYC | -111.620 | -4.896 | 60.568 | -14.995 | -70.943 |
| HON | -81.870 | -9.805 | 47.377 | -10.452 | -49.984 |

\*The binding energy is estimated from the sum of the van der Waal, electrostatic, polar solvation, and solvent accessible surface area (SASA) energies during the last 10 ns of the simulation.

Fig 6 illustrates the decomposition of the binding free energy of the strongest ligands (VER, ETI, CAR, HYD, CHL, ECL, CYC, and HON) on a per residue basis using MM/PBSA analysis. Residues Leu 128, Pro 132, and Val 155 significantly contributed to the negative binding free energy (Fig 6). Pro 132 and Leu 128 are also considered the gate residues of the NS2b-NS3 binding site and play a crucial role in inhibiting the replication process of the protein. Table on S3 Table can also be checked to determine which other residues may contribute on the binding energies per candidate ligands.

**Fig 6.**
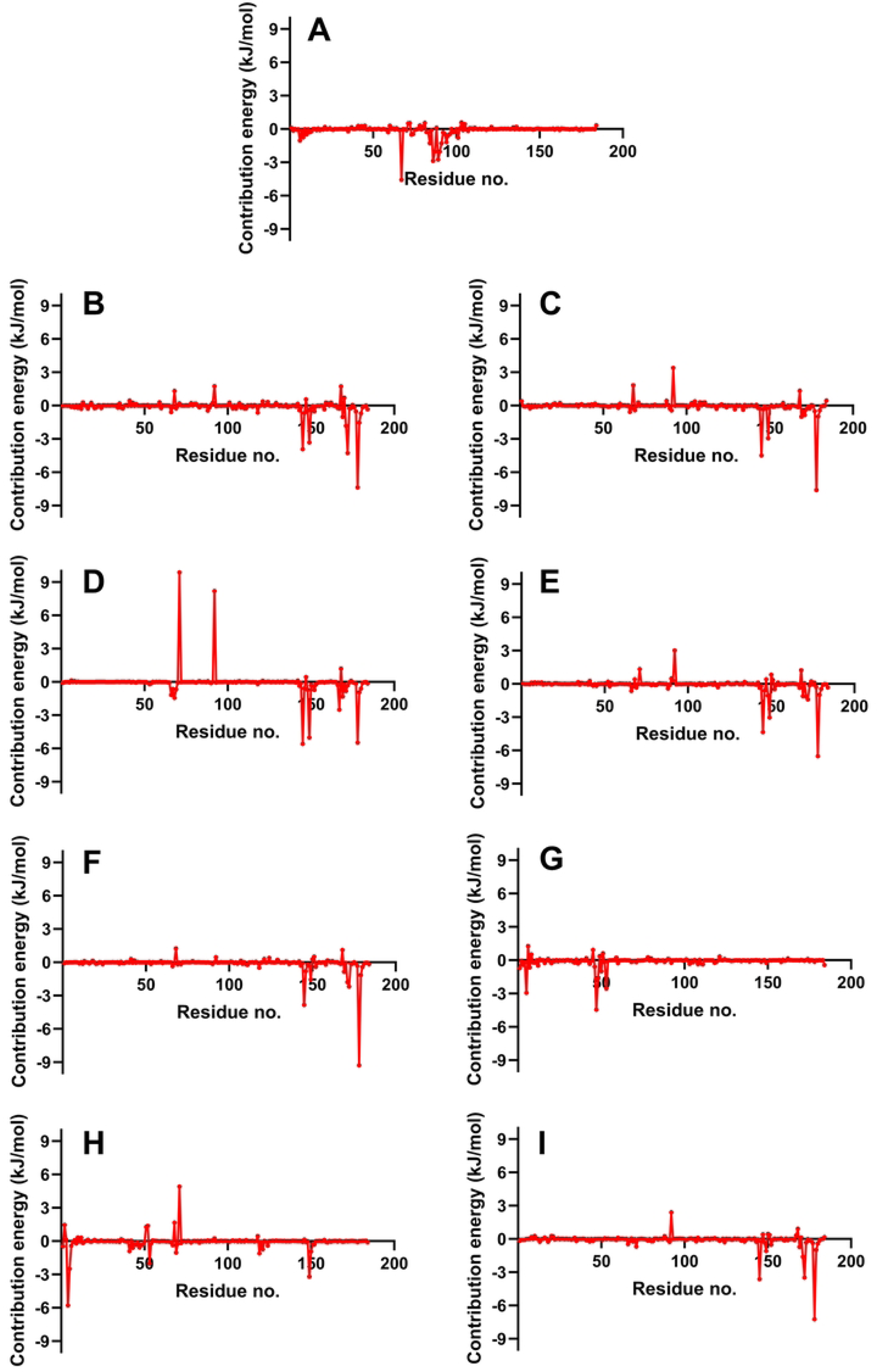
Binding energy decomposition on complexes with the highest MM/PBSA energies. Reference ligands: (A) NIN. Candidate ligands: (B) VER (C) ETI (D) CAR (E) HYD (F) CHL (G) ECL (H) CYC (I) HON

## Conclusion

In Philippine folkloric medicine, locals commonly used a decoction of *E. hirta, C. papaya,* or *M. charantia* as a treatment for dengue. To further examine the effectiveness of Philippine medicinal plants against dengue, the study evaluated 2,944 ligands from phytochemicals found naturally in the Philippines. ADMET analysis showed that 1,265 of the compounds are pharmacologically viable and safe for human consumption. Recent studies also showed that targeting the active site of NS2b-NS3 protease of DENV can inhibit viral replication and prevent viral infection. Idelalisib (IDE) and nintedanib (NIN), which are known drug molecules that can effectively bind to the active site of NS2b-NS3 protease, have been used as references in the study.

Molecular docking experiments were performed in the active site of the protease to find the top ten possible inhibitors from the 1,265 compounds. Results showed that all ten ligands (VER, ISO, ETI, TOM, CAR, HYD, CHL, and HON) have higher docking scores compared to the reference compounds IDE (−25.522 kJ/mol) and NIN (−29.790 kJ/mol). Several interactions between the ligands and active site residues contributed to the high docking scores, such as polar and hydrogen bonds near the active site residues Ser 135, Asp 75, and His 51.

The stability of the top ten complexes and references was evaluated by performing 100-ns MD simulations in triplicate. Analysis showed ISO and TOM, including the reference IDE were unstable based on their RMSD and RMSF trajectories. On the other hand, NIN and the other eight candidates have stable complex-RMSD and ligand-RMSD trajectories within the binding region, having an average RMSD of 3.2 Å on the last ten ns of MD simulation. MM/PBSA analysis further verified the binding affinities of the eight stable ligands against NS2b-NS3 protease, with NIN being the reference. Results showed the final six ligands with higher binding energies compared to NIN (−55.709 kJ/mol): VER (−80.682), ETI (−59.923), HYD (−61.951), CHL (−63.279), ECL (−56.932) and CYC (−70.943). CAR from *Carissa carandas* (carandas plum) and HON from *Agave americana* (century plant) can be potential dengue NS2b-NS3 protease inhibitors but had a lower MM/PBSA score than the reference drug.

Overall, the study identified six phytochemicals with high binding affinities against the NS2b-NS3 protease of the dengue virus: VER from *Veratrum mengtzeanum* (pimacao), ETI from *Lilion martagon* (Turk’s cap lily), HYD from *Eclipta prostrata* (false daisy), ECL from *Eclipta alba*, CHL from *Yucca gloriosa* (palm lily), and CYC from *Euphorbia hirta* (tawa-tawa). Results showed that the study might help explain the anecdotal efficacy of such folklore medicinal plants in the Philippines. More studies on contemporary pharmacological approaches, including the isolation of active compounds and elucidating the mode of action of antiviral activities, are warranted to validate the traditional claim overly.

## Acknowledgements

The authors would like to thank the Computing and Archiving Research Environment of the Department of Science and Technology – Advanced Science and Technology Institute (COARE-DOST-ASTI), the Engineering Research and Development for Technology of the Department of Science and Technology (ERDT-DOST), the University of the Philippines Engineering Research and Development Foundation, Inc (UP-ERDFI), and the University of the Philippines Diliman – Office of the Vice Chancellor for Research and Development (UP-OVCRD) for their support on this research.

## Supporting information captions

S1 Table. ADMET and toxicity results of the test ligands.

S2 Table. Docking results of test ligands and references on NS2b-NS3 protease (2FOM) using Autodock 4.2.

S3 Table. Decomposition of binding free energy on a per residue basis of the complex with strongest MM/PBSA energies

